# PICK1 links KIBRA and AMPA receptors in coiled-coil-driven supramolecular complexes

**DOI:** 10.1101/2024.03.12.584494

**Authors:** Xin Shao, Lenora Volk

## Abstract

The human memory-associated protein KIBRA regulates synaptic plasticity and trafficking of AMPA-type glutamate receptors, and is implicated in multiple neuropsychiatric and cognitive disorders. How KIBRA forms complexes with and regulates AMPA receptors remains unclear. Here, we show that KIBRA does not interact directly with the AMPA receptor subunit GluA2, but that PICK1, a key regulator of AMPA receptor trafficking, can serve as a bridge between KIBRA and GluA2. We identified structural determinants of KIBRA-PICK1-AMPAR complexes by investigating interactions and cellular expression patterns of different combinations of KIBRA and PICK1 domain mutants. We find that the PICK1 BAR domain, a coiled-coil structure, is sufficient for interaction with KIBRA, whereas mutation of the BAR domain disrupts KIBRA-PICK1-GluA2 complex formation. In addition, KIBRA recruits PICK1 into large supramolecular complexes, a process which requires KIBRA coiled-coil domains. These findings reveal molecular mechanisms by which KIBRA can organize key synaptic signaling complexes.

## Introduction

The *KIBRA* gene (enriched in KIdney and BRAin) is associated with human memory performance (1–13), and *Kibra* disruption impairs learning and memory in rodents (14–17). *KIBRA* polymorphisms and gene expression also associate with disorders of complex brain function including schizophrenia (18), autism spectrum disorder (19), and Tourette syndrome (20). KIBRA is a scaffolding protein enriched at excitatory synapses (14, 21), with a large proportion of the KIBRA interactome also implicated in neuropsychiatric disorders (18, 22–27). Additionally, decreased KIBRA protein expression in the brain correlates with tauopathy-related cognitive impairment in humans (28, 29).

The involvement of KIBRA in the mechanisms underlying learning and memory remains incompletely understood. However, KIBRA has been shown to regulate synaptic plasticity, a cellular correlate of memory (30, 31), in multiple model organisms (14–17, 32, 33). KIBRA is also necessary for experience-induced plasticity of *in vivo* hippocampal and cortical circuit dynamics (34). Moreover, multiple studies have demonstrated that KIBRA can regulate the trafficking and expression levels AMPA receptors, ionotropic glutamate receptors primarily responsible for fast excitatory neurotransmission in the central nervous system (14, 15, 17). As AMPA receptor trafficking is a highly conserved expression mechanism underlying many forms of excitatory synaptic plasticity (35, 36), KIBRA’s influence on AMPA receptor trafficking is hypothesized to underlie its regulatory function in synaptic plasticity. While the specific cellular and molecular mechanisms underlying KIBRA’s regulation of AMPA receptor trafficking remain unclear, experimental evidence indicates that KIBRA can form a complex with native AMPA receptors and regulators of AMPA receptor trafficking including GRIP1, NSF, Sec8, and PICK1 *in vivo* (14). Additionally, there is evidence supporting direct interaction between KIBRA and PICK1 (14). PICK1 interacts with the GluA2 subunit of AMPARs, and regulates synaptic plasticity and memory (37–39). It is not known if KIBRA binds GluA2 directly or if PICK1 can serve as a bridge between KIBRA and GluA2. Additionally, the molecular determinants regulating PICK1-KIBRA interactions are unknown.

KIBRA is a scaffolding protein containing multiple protein-protein and protein-lipid interaction domains (40, 41) (see Fig. 6F): two N-terminal WW domains (protein-interaction domains that bind PPxY motifs and other proline-rich regions) (42–44), central and C-terminal coiled-coil domains (CC domains mediate homotypic or heterotypic association with other coiled coils and facilitate oligomerization) (45, 46), a C2-like domain (mediating lipid binding, likely in a Ca^2+^-independent manner) (40, 47), a glutamate-rich region, an atypical PKC-binding region (αBR) and a C-terminal type III PDZ (Postsynaptic Density-95/Discs large/Zonula occludens-1) ligand (mediating interaction with proteins containing PDZ domains) (48, 49). WW domain/PPxY interactions are ubiquitous among members of the HIPPO signaling pathway, which plays a conserved role regulating cell growth, division, and organ size (42–44). Interestingly, WW domain mutants that disrupt KIBRA interaction with HIPPO signaling molecules exhibit stronger interaction with synaptic and AMPAR regulatory protein complexes, and hippocampal overexpresion of these KIBRA WW-domain mutants improves memory (50).

**Figure 1.**
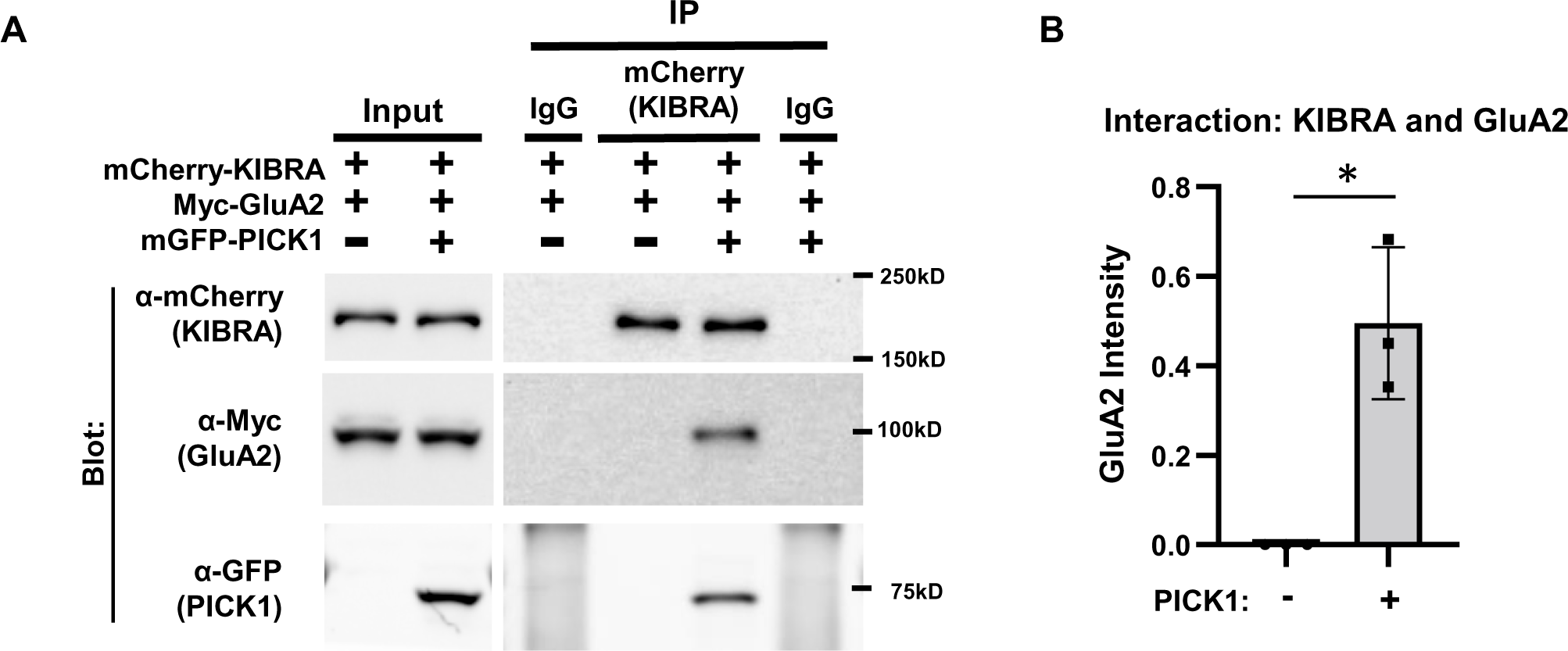
PICK1 mediates interaction between KIBRA and GluA2. **(A)** mCherry-KIBRA and Myc-GluA2 were transfected into HEK293T cells with either mGFP or mGFP-PICK1. mCherry antibody was used to immunoprecipitate mCherry-KIBRA, and co-precipitated Myc-GluA2 and mGFP-PICK1 were detected by immunoblot. **(B)** Quantification of 3 biological replicates showed that GluA2 was only coprecipitated with KIBRA in the presence of PICK1. GluA2 signal without PICK1: 0, with PICK1: 0.50 ± 0.17; n=3; Paired t-test, *p<0.05. Data reported as mean ± SD.

**Figure 2.**
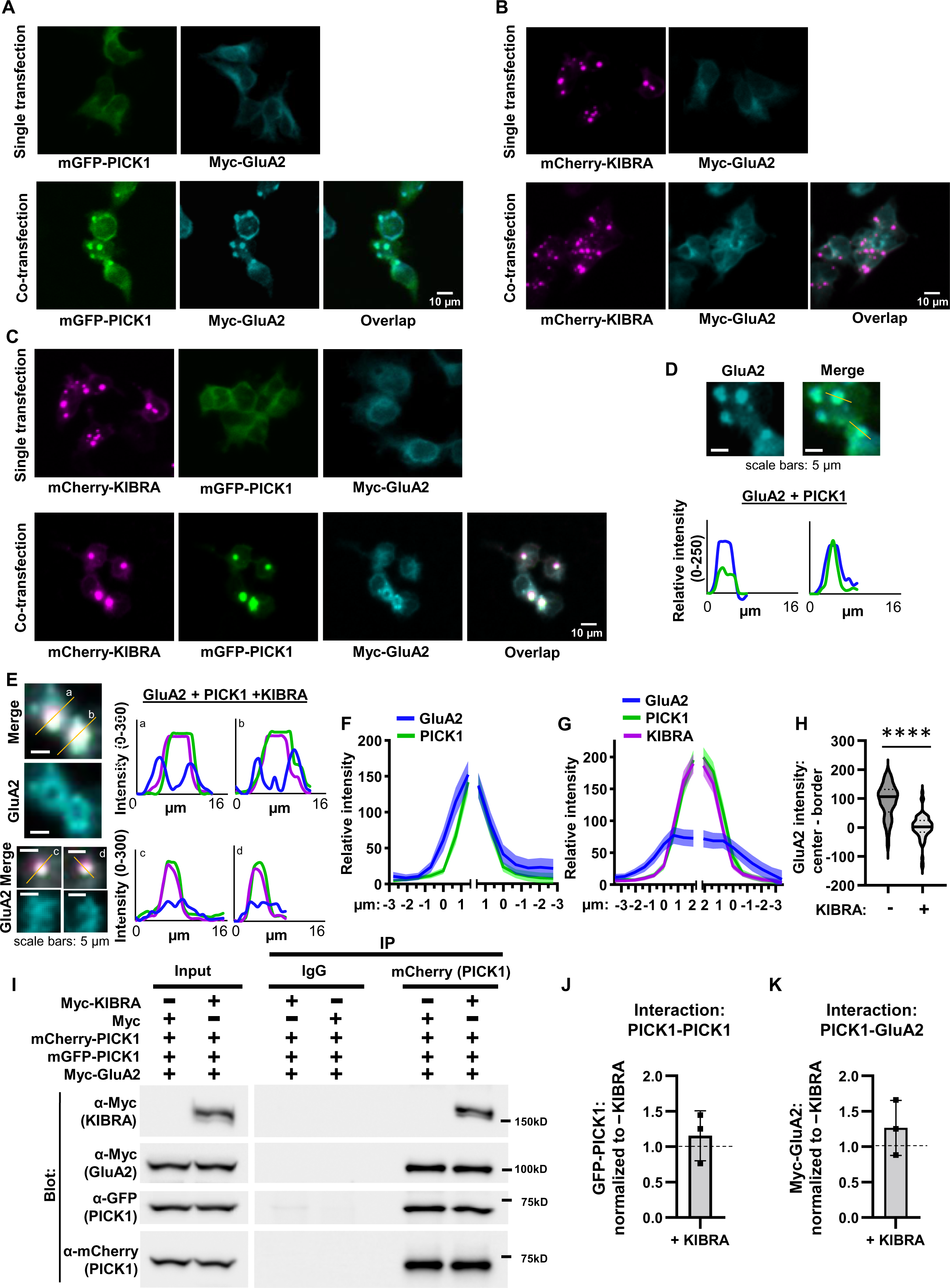
KIBRA promotes PICK1 puncta formation but alters PICK1/GluA2 co-clusters independent of PICK1 dimerization or PICK1 interaction with GluA2. (**A)** mGFP-PICK1 and Myc-GluA2 were transfected into HEK293T cells. Singly transfected PICK1 and GluA2 show diffuse localization (top) whereas co-transfected PICK1 and GluA2 form clusters (bottom). **(B)** mCherry-KIBRA and Myc-GluA2 were transfected into HEK293T cells. KIBRA forms puncta whether singly transfected (top) or co-transfected with GluA2 (bottom), but does not affect the diffuse localization of GluA2. **(C)** mCherry-KIBRA, mGFP-PICK1 and Myc-GluA2 were transfected into HEK293T cells. Co-transfected KIBRA alters PICK1 and GluA2 co-clusters. **(D)** Example line scans of GluA2 and PICK1 in the indicated clusters from panel A. **(E)** Example line scans of GluA2, KIBRA, and PICK1 signals from the indicated clusters in panel C. **(F, G)** Summary of line scan quantification for HEK293T cells transfected with GluA2 and PICK1 (F) or GluA2, PICK1, and KIBRA (G). Data are plotted as mean ± 95% CI. Signals were aligned to the edges (x = 0) of each KIBRA/PICK1 (F) or PICK1 (G) cluster. **(H)** GluA2 signal intensity at the border of each cluster was subtracted from the signal at the cluster center. Dark lines on violin plot denote median, dashed lines indicate quartiles. Mann-Whitney test, p < 0.0001; median and 95% CI limits: PICK1 +GluA2: = 106.3, 79.88 to 121.2, PICK1 + GluA2 + KIBRA =2.447, −6.371 to 12.5. n = 36 cells for PICK1/GluA2-expressing cells and 70 cells for PICK1/GluA2/KIBRA-expressing cells, each from three biological replicates. Signals in D-H were measured following local background subtraction. **(I)** mGFP/mCherry-PICK1 and Myc-GluA2 were transfected into HEK293T cells with Myc or Myc-KIBRA. Anti-mCherry antibodies were used to immunoprecipitate mCherry-PICK1, and co-precipitated Myc-KIBRA, Myc-GluA2 and mGFP-PICK1 were detected by immunoblot using mGFP or Myc antibodies. **(J, K)** PICK1 homodimerization (J, KIBRA+ normalized to KIBRA negative sample within the same gel: 1.15 ± 0.35; p>0.05) and PICK1-GluA2 interaction (K, KIBRA+, normalized to KIBRA-negative sample within the same gel: 1.26 ± 0.39; p>0.05) were not affected by KIBRA expression, quantified from 3 biological replicates. One sample t-test vs. 1. Data reported as mean ± SD unless otherwise noted. Scale bars for A, B, C = 10 µm, scale bars in D and E = 5 µm.

**Figure 3.**
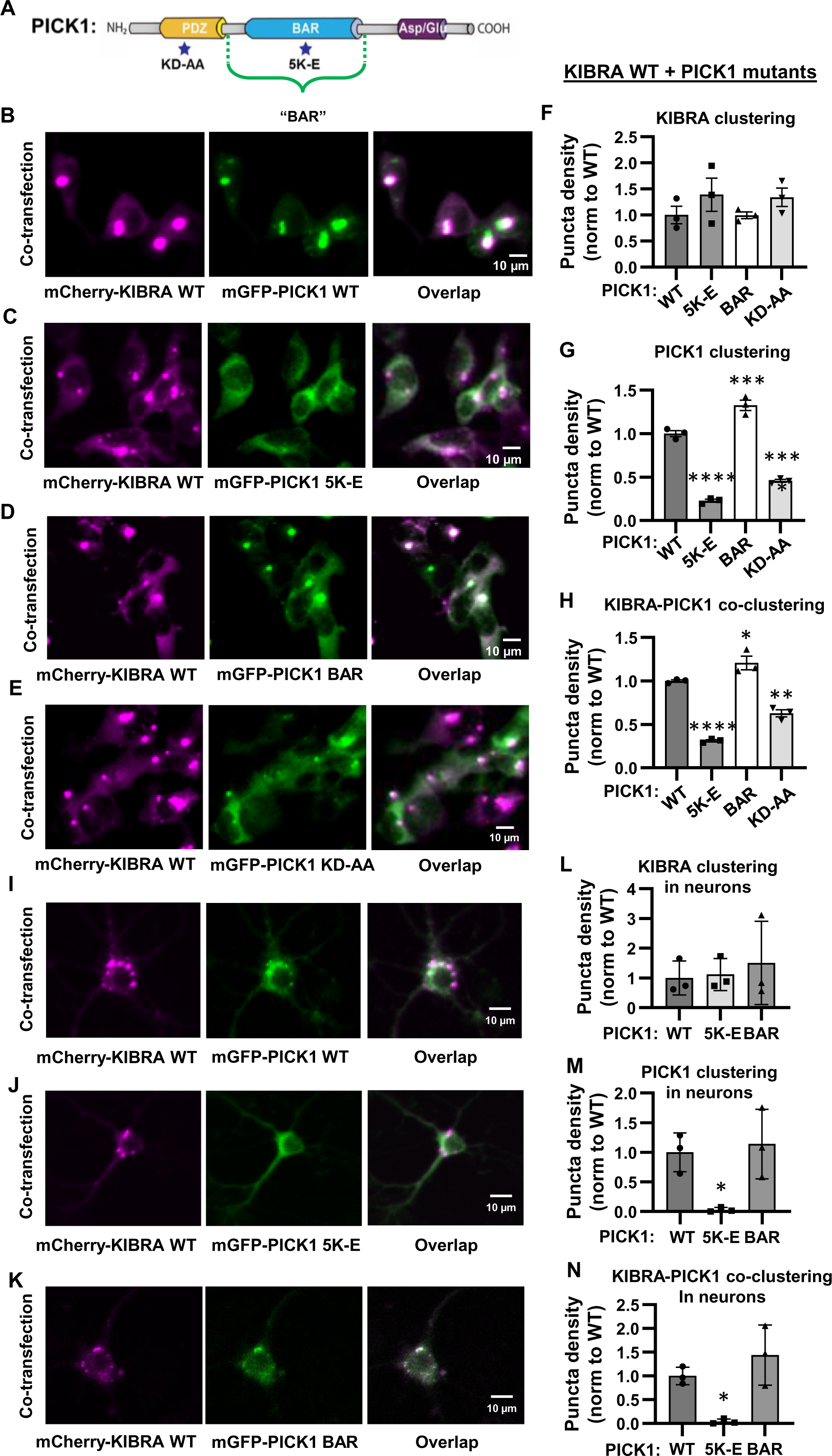
PICK1 BAR domain regulates the ability of PICK1 to co-aggregate with KIBRA. **(A)** PICK1 domain schematic with domain mutations. **(B-E)** Representative images of HEK-393T cells co-expressing KIBRA WT with (**B)** PICK1 WT**, (C)** PICK1 BAR domain mutant, 5K-E, **(D)** isolated PICK1 BAR domain, or **(E)** PICK1 PDZ-domain mutant, KD-AA. PICK1 WT and isolated BAR domain form clusters with KIBRA (B,D). Point mutants that disrupt BAR domain (5K-E) or PDZ domain (KD-AA) function result in diffuse PICK1 localization without affecting KIBRA clustering in HEK293T cells (C,E). **(F-H)** Quantification of 3 biological replicates from conditions shown in B-E; **(F)** Density of WT KIBRA puncta when co-expressed with PICK1 WT (1.00 ± 0.29), PICK1 5K-E (1.39 ± 0.55), PICK1 BAR (0.99 ± 0.11), PICK1 KD-AA (1.34±0.31). One-way ANOVA, p = 0.4051, Dunnett’s multiple comparisons test vs. WT, p > 0.05 for all PICK1 mutants. **(G)** Density of PICK1 WT and mutant puncta when co-expressed with KIBRA WT; PICK1 WT (1.00 ± 0.05), PICK1 5K-E (0.23 ± 0.03), PICK1 BAR (1.33 ± 0.10), PICK1 KD-AA (0.46 ± 0.03). One-way ANOVA, p < 0.0001, Dunnett’s multiple comparisons test vs. WT ***p < 0.001, ****p < 0.0001. **(H)** Density of co-localized KIBRA WT and PICK1 WT (1.00 ± 0.02), PICK1 5K-E (0.31 ± 0.02), PICK1 BAR (1.21 ± 0.14), and PICK1 KD-AA (0.63 ± 0.07). One-way ANOVA p < 0.0001, Dunnett’s multiple comparisons test vs. WT *p < 0.05, **p < 0.01, ****p < 0.0001. **(I-K)** WT KIBRA with isolated PICK1 BAR domain or BAR domain mutant were expressed in hippocampal neurons cultured from PICK1 KO mice. PICK1 WT and isolated BAR domain form clusters with KIBRA while PICK1 5K-E shows diffuse localization. **(L-N)** Quantification of 3 biological replicates from conditions shown in I, J, and K; **(L)** density of WT KIBRA puncta when co-expressed with PICK1 WT (1.00 ± 0.57), PICK1 5K-E (1.12 ± 0.54), PICK1 BAR (1.50 ± 1.34). One-way ANOVA, p = 0.7903, Dunnett’s multiple comparisons test vs. WT, p > 0.05 for all PICK1 mutants. **(M)** Density of PICK1 WT and mutant puncta when co-expressed with KIBRA WT; PICK1 WT (1.00 ± 0.33), PICK1 5K-E (0.03 ± 0.04), PICK1 BAR (1.14 ± 0.59). One-way ANOVA, p = 0.0253, Dunnett’s multiple comparisons test vs. WT *p < 0.05. **(N)** Density of co-localized KIBRA WT with PICK1 WT (1.00 ± 0.18), PICK1 5K-E (0.04 ± 0.05), PICK1 BAR (1.44 ± 0.63). One-way ANOVA, p = 0.0108, Dunnett’s multiple comparisons test vs. WT *p < 0.05. Data reported as mean ± SD. Scale bars in B-E and I-K: 10 µm. For quantification in F-H and L-N, all values were normalized to the average of the three WT biological replicates (independent cultures).

**Figure 4.**
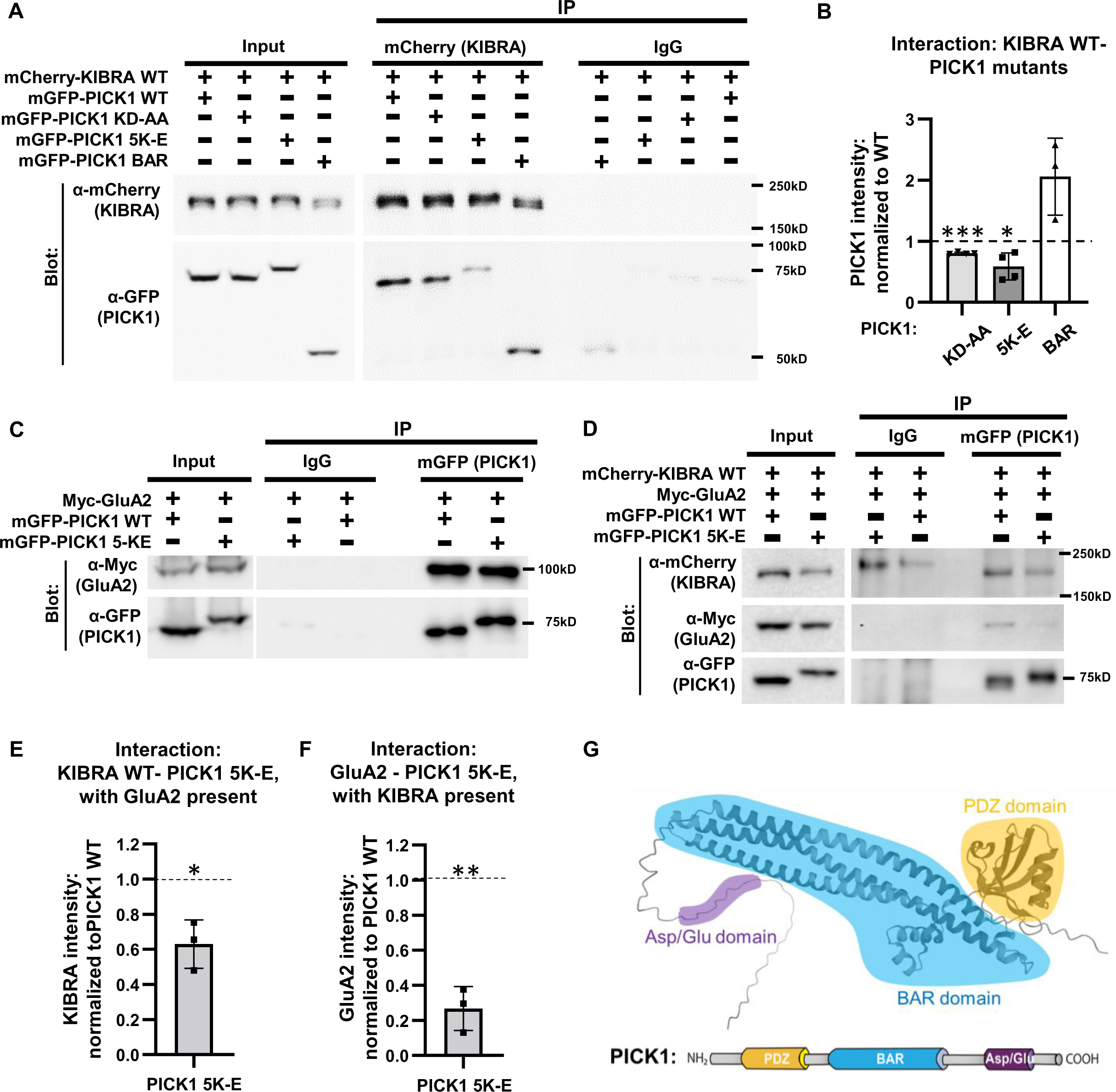
PICK1 BAR domain facilitates the interaction between KIBRA and PICK1. **(A)** mGFP-PICK1 WT, KD-AA, 5K-E or BAR was transfected with mCherry-KIBRA WT into HEK293T cells. mCherry-KIBRA was immunoprecipitated and co-precipitated mGFP-PICK1 variants were detected by immunoblot using anti-GFP antibodies. **(B)** Quantification of 4 biological replicates for PICK1 WT, KD-AA and 5K-E and 3 biological replicates for PICK1 BAR as shown in A; PICK1 KD-AA or 5K-E impair interaction with KIBRA compared with PICK1 WT (PICK1 pull down normalized to input and same-gel WT PICK1; PICK1 KD-AA: 0.80 ± 0.02, **p<0.01; PICK1 5K-E: 0.59 ± 0.22, *p<0.05; PICK1 BAR: 2.06 ± 0.63, n=3; p = 0.1008, one sample t-test vs. 1). **(C)** mGFP-PICK1 WT or 5K-E and Myc-GluA2 were transfected into HEK293T cells. Immunoprecipitation of PICK1 was performed using a GFP antibody, and the co-precipitated GluA2 was detected using a Myc antibody. GluA2 showed equivalent interaction with mGFP-PICK1 WT and 5K-E. **(D)** mGFP-PICK1 WT or 5K-E was co-transfected with mCherry-KIBRA and Myc-GluA2 into HEK293T cells. mGFP-PICK1 was immunoprecipitated and co-precipitated mCherry-KIBRA and Myc-GluA2 were detected by immunoblot. **(E, F)** Quantification of 3 biological replicates of conditions shown in D; When PICK1, KIBRA and GluA2 are all present, mutation of the PICK1 BAR domain (PICK1 5K-E) decreases interaction with both KIBRA and GluA2. (All bands were normalized to input. KIBRA signal with PICK1 5K-E normalized to +PICK1 WT: 0.63 ± 0.14, *p<0.05; GluA2 signal with PICK1 5K-E normalized to +PICK1 WT: 0.27 ± 0.13, **p<0.01, one sample t-test.) **(G)** Structure and domains of PICK1. Predicted structure modified from AlphaFold Protein Structure Database. Data reported as mean ± SD.

**Figure 5.**
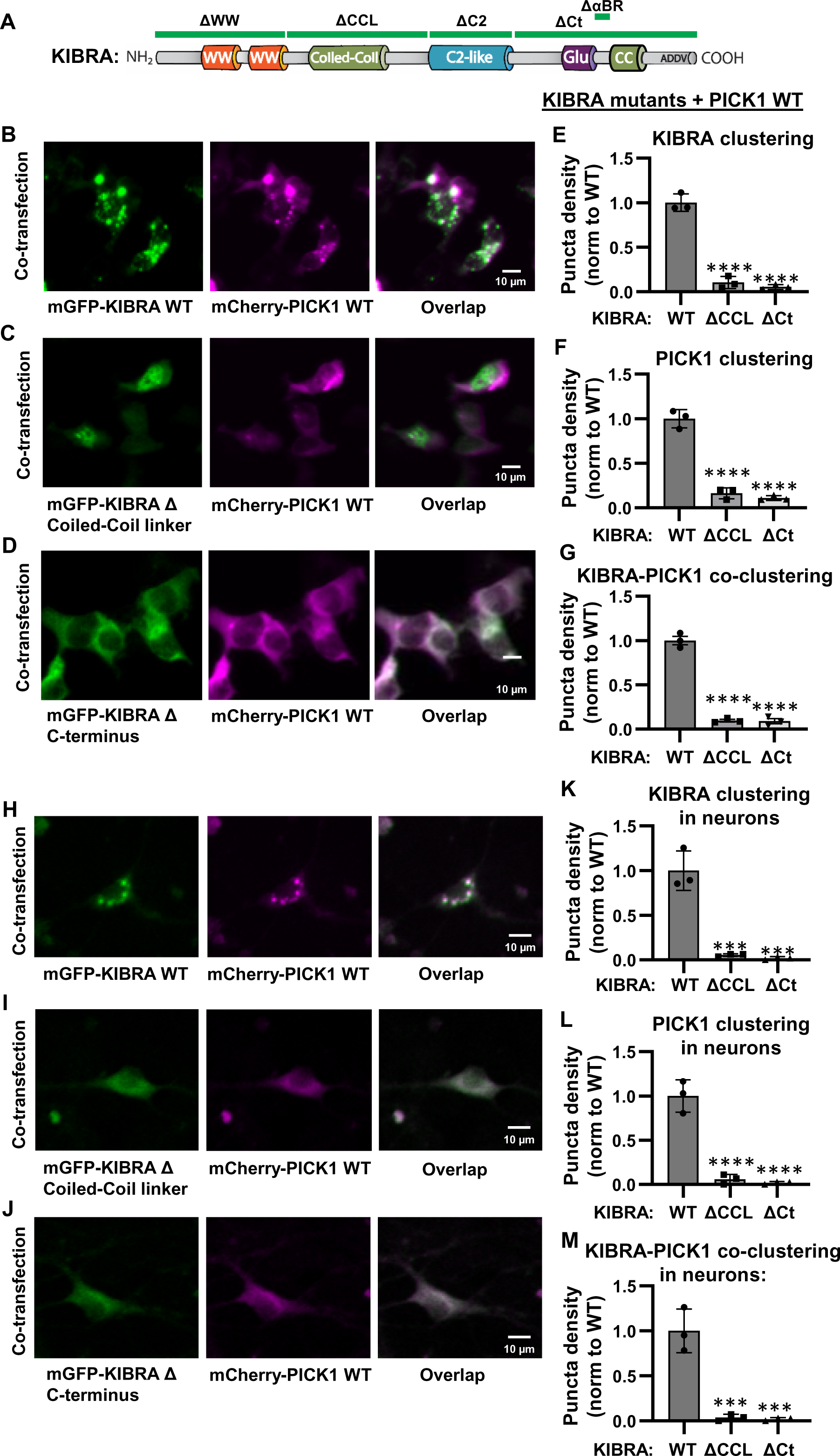
KIBRA coiled-coil-containing domains contribute to KIBRA clustering and subsequent PICK1 co-aggregation. **(A)** KIBRA domain schematic with the location of domain truncations noted. **(B-D)** KIBRA WT clusters with PICK1 WT while KIBRA mutants missing the coiled-coil linker or C-terminal regions (ΔCCL and ΔCt, respectively) show diffuse localization and fail to cluster PICK1 in HEK293T cells. **(E-G)** Quantification of 3 biological replicates for conditions shown in B, C, and D. **(E)** KIBRA puncta density for KIBRA WT (1.00 ± 0.10), KIRBA ΔCCL (0.10 ± 0.07), and KIBRA ΔCt (0.05 ± 0.03). One-way ANOVA, p < 0.0001, Dunnett’s multiple comparisons test vs. WT, ****p < 0.0001. **(F)** PICK1 cluster density when co-expressed with KIBRA WT (1.00 ± 0.10), ΔCCL (0.16 ± 0.06), or ΔCt (0.11 ± 0.03). One-way ANOVA p < 0.0001, Dunnett’s multiple comparisons test vs. WT, ****p < 0.0001. **(G)** PICK1-KIBRA co-cluster density for KIBRA WT (1.00 ± 0.08), ΔCCL (0.10 ± 0.02), or ΔCt (0.09 ± 0.05). One-way ANOVA, p < 0.0001, Dunnett’s multiple comparisons test vs. WT, ****p < 0.0001. **(H-J)** KIBRA WT, ΔCCL or ΔCt were expressed with WT PICK1 in neurons lacking endogenous KIBRA (cultures prepared from KIBRA KO mice). WT KIBRA clusters with PICK1 in mouse hippocampal neurons while KIBRA ΔCCL and ΔCt show diffuse localization and fail to cluster PICK1. **(K-M)** Quantification of 3 biological replicates for conditions shown in H, I and J. **(K)** KIBRA puncta density for KIBRA WT (1.00 ± 0.22), KIRBA ΔCCL (0.05 ± 0.01), and KIBRA ΔCt (0.02 ± 0.02). One-way ANOVA, p = 0.0001, Dunnett’s multiple comparisons test vs. WT, ***p < 0.001. **(L)** PICK1 cluster density when co-expressed with KIBRA WT (1.00 ± 0.18), ΔCCL (0.06 ± 0.06), or ΔCt (0.18 ± 0.02). One-way ANOVA p < 0.0001, Dunnett’s multiple comparisons test vs. WT, ****p < 0.0001. **(M)** PICK1-KIBRA co-cluster density for KIBRA WT (1.00 ± 0.24), ΔCCL (0.04 ± 0.04), or ΔCt (0.02 ± 0.02). One-way ANOVA, p = 0.0002, Dunnett’s multiple comparisons test vs. WT, ***p < 0.001.Data reported as mean ± SD. Scale bar for B-D and H-J: 10 µm. For quantification in E-G and K-M, all values were normalized to the average of the three WT cultures.

**Figure 6.**
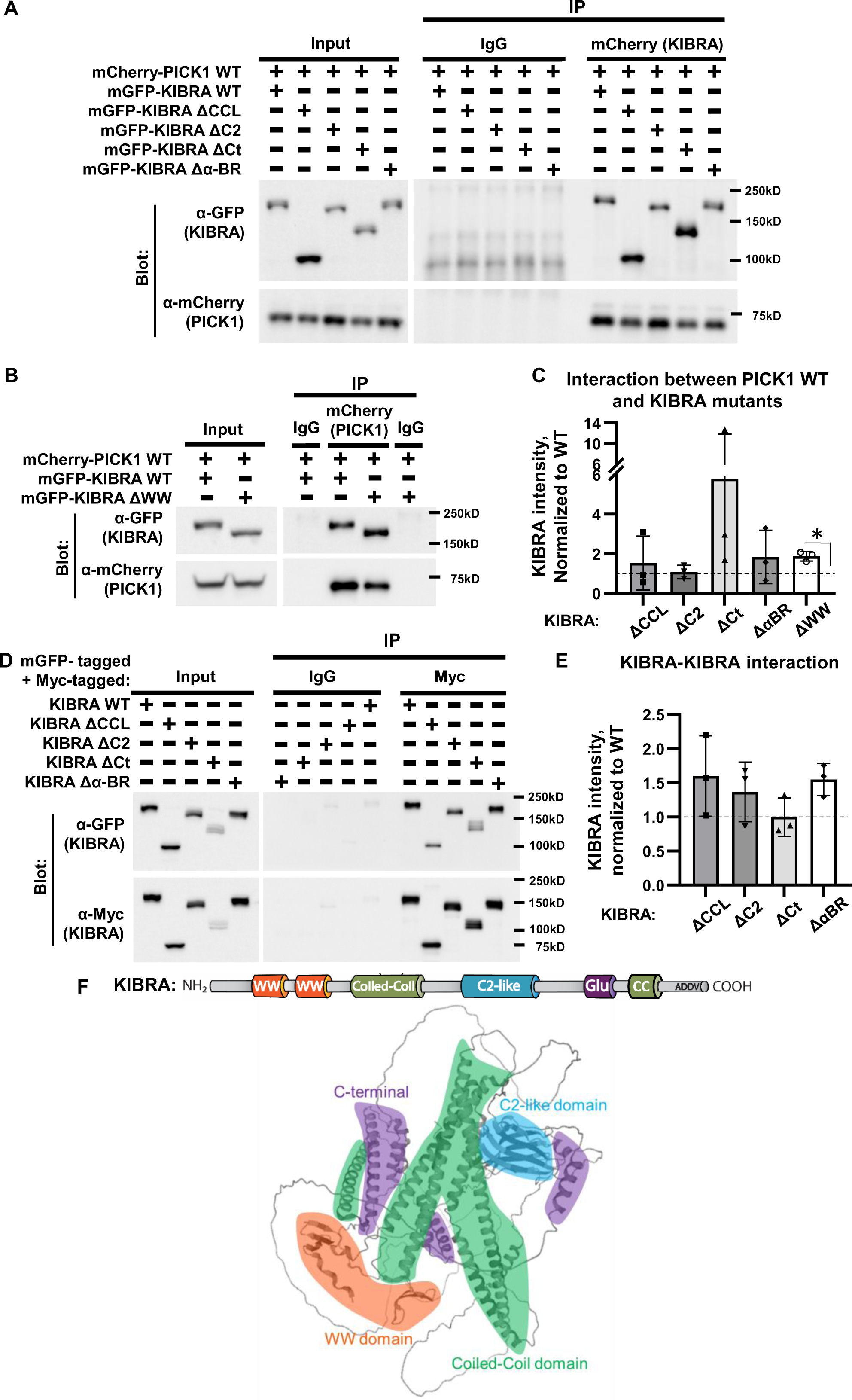
No single domain of KIBRA is essential for interaction between KIBRA and PICK1 or for KIBRA-KIBRA dimerization. **(A, B)** mGFP-KIBRA WT or mutants missing the coiled-coil linker, C2 domain, C-terminal region, aPKC binding region, or WW domain (ΔCCL, ΔC2, ΔCt, ΔαBR, ΔWW, respectively) were transfected with mCherry-PICK1 WT into HEK293T cells. Anti-mCherry antibodies were used to immunoprecipitate PICK1, and co-precipitated mGFP-KIBRA variants were detected by immunoblot for GFP. **(C)** Quantification of 3 biological replicates under conditions shown in A and B. None of the KIBRA mutants disrupt interaction between KIBRA and PICK1 (values normalized to WT sample from the same blot; ΔCCL 1.53 ± 1.34, ΔC2 1.08 ± 0.24, ΔCt 5.79 ± 6.01, ΔαBR 1.84 ± 1.34, one-sample t-test vs 1, p > 0.10). KIBRA ΔWW shows increased interaction with PICK1 (ΔWW 1.87 ± 0.24, one-sample t-test vs 1, p < 0.05). **(D)** Myc-and GFP-KIBRA WT, ΔCCL, ΔC2, ΔCt, or ΔαBR were transfected into HEK293T cells. Myc-KIBRA was immunoprecipitated and co-precipitated mGFP-KIBRA was detected by immunoblot. **(E)** Quantification of 3 biological replicates under conditions shown in D. No single KIBRA mutant prevents homodimerization (values normalized to WT sample from the same blot; ΔCCL 1.60 ± 0.59, ΔC2 1.36 ± 0.43, ΔCt 1.00 ± 0.28, ΔαBR 1.55 ± 0.23, one-sample t-test vs 1, p > 0.05. **(F)** Structure and domains of KIBRA. Predicted structure modified from AlphaFold Protein Structure Database. Data reported as mean ± SD.

PICK1 is unusual in that it contains both a PDZ domain and a BAR (Bin/Amphiphysin/RVS) domain (51). PDZ domains support protein-protein interactions with proteins containing C-terminal PDZ ligands, while BAR domains bind lipids, sense/regulate membrane curvature, and are commonly found in proteins involved in membrane trafficking (51, 52). The PICK1 PDZ domain preferentially binds type I and II PDZ ligands, among which are the GluA2 and GluA3 AMPAR subunits (51, 53). PICK1 BAR domain function is required for membrane trafficking of AMPARs (54, 55). BAR domains are composed of alpha-helical bundles that can engage in coiled-coil interactions with other proteins, including promotion of PICK1 dimerization (54, 56–60). Notably, the PICK1 PDZ and BAR domains can interact with each other, promoting a closed conformation that disrupts some BAR domain interactions. Binding a PDZ ligand (e.g. GluA2) facilitates membrane localization and shifts PICK1 to an open conformation, promoting BAR domain activation (56, 58, 61–63).

To gain insight into mechanisms by which KIBRA regulates AMPAR complexes, we investigated the molecular basis of PICK1-KIBRA interactions by introducing mutations within the major functional domains of KIBRA and PICK1. Investigation of protein interactions and cellular expression patterns of protein complexes revealed that KIBRA does not bind GluA2 directly and that PICK1 can serve as a bridge between KIBRA and GluA2. Our data further indicate that the PICK1 BAR domain regulates interaction with KIBRA, and that multiple domains of KIBRA can interact with PICK1. Additionally, we show that KIBRA initiates coiled-coil domain-dependent formation of large, supramolecular clusters with PICK1.

## Results

### PICK1 mediates interaction between KIBRA and GluA2

While KIBRA plays a crucial role in regulating AMPAR trafficking, activity-induced increases in AMPAR expression, and synaptic plasticity (14, 15, 17, 50), how KIBRA complexes with AMPARs remains an open question. KIBRA interacts directly with the AMPAR-regulatory protein PICK1, which binds the GluA2 subunit of AMPARs through PDZ domain (PICK1)/PDZ ligand (GluA2) interactions (37, 64). To test the hypothesis that PICK1 facilitates KIBRA-AMPAR complex formation, we performed Co-IP (Co-Immunoprecipitation) experiments in HEK293T cells transfected with mCherry-KIBRA, Myc-GluA2, and either mGPF or mGFP-PICK1. Following immunoprecipitation of mCherry-KIBRA, co-precipitated GluA2 was detected using a Myc antibody. We found that Myc-GluA2 was only detected in the presence of co-expressed mGFP-PICK1, but not with mGFP alone (Fig. 1A, B), indicating that KIBRA does not directly interact with GluA2, and that PICK1 can serve as a bridge between KIBRA and GluA2.

To further investigate the function of KIBRA in AMPAR complexes, we examined the expression pattern of GluA2 alone or co-expressed with PICK1 or KIBRA. Consistent with prior reports (14, 37, 64), PICK1 and GluA2 show diffuse cytoplasmic localization when expressed alone, but form co-localized clusters when expressed together (Fig 2A,D,F). KIBRA forms large clusters when expressed alone (Fig. 2B) (14). We observed no co-clustering between co-expressed KIBRA and GluA2 (Fig. 2B), consistent with results from our co-immunoprecipitation experiments indicating that KIBRA does not directly interact with GluA2.

### KIBRA promotes co-aggregation of PICK1 and modulates PICK1-GluA2 clustering

It is well established that PICK1 plays a crucial role in the trafficking of GluA2-containing AMPA receptors (56). PICK1 clusters GluA2 in heterologous mammalian cells (Fig 2A, G) (37) and regulates AMPAR trafficking in neurons (38, 58, 61, 65, 66). Because KIBRA clusters PICK1 in heterologous mammalian cells (Fig 3A, 5A) (14) and also regulates AMPAR trafficking in neurons (14, 17), we predicted that KIBRA, PICK1, and GluA2 would form overlapping tripartite clusters when co-expressed. Surprisingly, while KIBRA and PICK1 still formed large overlapping clusters in the presence of GluA2, GluA2 concentrations were not precisely localized with PICK1/KIBRA clusters (Fig. 2C). We observed rings of GluA2 concentration surrounding some PICK1/KIBRA clusters (Fig. 2C, E, G). Line scans though the center of PICK1/KIBRA clusters revealed that while GluA2 does concentrate with PICK1/KIBRA clusters (Fig. 2G, GluA2 signal at the edge of PICK1/KIBRA clusters is significantly elevated compared to cellular background. Border mean ± SEM= 107.3 ± 5.8, background = 71.6 ± 5.0, Wilcoxin test p < 0.0001, n = 70 cells from 3 biological replicates), the precise spatial co-localization of GluA2 with PICK1 is altered by the presence of KIBRA (Fig. 2 F, G, H). Intriguingly, the spatial organization of tripartite clusters, with KIBRA and PICK1 at the core surrounded by GluA2, is strikingly reminiscent of liquid-liquid phase separated (LLPS) postsynaptic density complexes that have been suggested to segregate different glutamate receptors and their scaffolds into adjacent nanodomains (67). Consistent with this observation, KIBRA was recently shown to initiate formation of LLPS condensates to regulate HIPPO signaling in non-neuronal cells (68, 69).

To determine if the redistribution of PICK1-GluA2 complexes in the presence of KIBRA results from altered PICK1-GluA2 interaction or altered PICK1 dimerization, we performed co-immunoprecipitations from HEK293T cells transfected with mGFP-PICK1, mCherry-PICK1, and Myc-GluA2 along with either Myc or Myc-KIBRA (Fig. 2I-K). We find that KIBRA does not affect PICK1-GluA2 or PICK1-PICK1 interaction, suggesting that the reorganization of PICK1 and GluA2 into LLPS-like clusters by KIBRA is not due to altered interaction between PICK1 and GluA2 or disruption of PICK1 dimerization.

### The BAR domain of PICK1 mediates co-aggregation of PICK1 with KIBRA

PICK1 consists of two major functional domains: a BAR domain which facilitates PICK1 interaction with phospholipids and actin, and a PDZ domain responsible for interaction with proteins containing C-terminal PDZ ligands such as GluA2 (37, 54, 64). The cooperative action of the PDZ and BAR domains is vital for regulating AMPA receptor trafficking (56). While PICK1 and KIBRA have been shown to interact in heterologous cells and *in vivo* (14), the nature of this interaction is unknown. To investigate the specific domains that regulate PICK1-KIBRA complex formation, we generated multiple PICK1 mutants that either1) disrupt PDZ domain interaction with proteins containing PDZ ligands (KD-AA) (37, 70–75), 2) disrupt BAR domain interactions (5K-E) (54), or 3) truncate PICK1 to contain only the isolated BAR domain (Fig. 3A). Co-localized clustering between PICK1 and KIBRA was observed as expected upon transfection of mCherry-KIBRA WT with mGFP-PICK1 WT (Fig. 3B) (14). The relative density of KIBRA puncta remained unchanged when co-expressed with PICK1 mutants or truncations (Fig 3C-F), consistent with data demonstrating that KIBRA can form clusters when expressed alone (Fig2B, C) (14, 68). However, PICK1 clustering and the formation of PICK1-KIBRA co-clusters was reduced when KIBRA WT was co-transfected with the PICK1 BAR domain mutant (5K-E) (Fig. 3C), whereas co-transfection of KIBRA WT with the isolated PICK1 BAR domain still led to the formation of PICK1 co-clusters with KIBRA (Fig. 3D). Quantitative analysis showed that BAR domain mutation decreased the relative density of PICK1 clusters and the overlap between KIBRA and PICK1 puncta (Fig. 3G, H). The isolated PICK1 BAR domain was more effective than WT PICK1 at promoting PICK1 clustering with KIBRA (Fig. 3F, G),suggesting that the BAR domain is sufficient to mediate KIBRA-induced clustering of PICK1, and that other PICK1 domains may serve as regulatory elements influencing this interaction. While KIBRA does contain a PDZ ligand, it is a type III sequence, whereas PICK1 preferentially binds type I and II PDZ ligands, with a higher affinity for type II ligands (51). Additionally, the KIBRA fragment identified in the original yeast two-hybrid screen for PICK1 interactors did not contain the PDZ ligand (14). However, PICK1 BAR and PDZ domains can form intramolecular interactions, promoting a closed conformation that inhibits some intermolecular BAR-domain interactions (56, 58, 61, 62). Consistent with this model, we observe that disrupting PICK1 PDZ domain interactions (PICK1-KD-AA) also decrease KIBRA-induced PICK1 cluster formation and co-localization of PICK1 and KIBRA puncta (Fig. 3 E, G, H).

We further validated the importance of the PICK1 BAR domain for KIBRA-induced PICK1 clustering in hippocampal neurons. We expressed PICK1 BAR domain mutants or isolated BAR domain in neurons cultured from PICK1 knockout mice to avoid confounds from endogenous WT PICK1 expression. At DIV10, neurons were infected with lentivirus expressing mCherry-KIBRA WT and mGFP-PICK1 WT, 5K-E, or BAR, and imaging was conducted at DIV14. Consistent with the results obtained in HEK293T cells, PICK1 WT and isolated PICK1 BAR domain formed co-clusters with KIBRA WT in neurons (Fig. 3I, K, M, N). However, PICK1 5K-E Bar domain mutant protein failed to form co-clusters with KIBRA (Fig. 3J, M, N). The consistent results obtained in both neurons and HEK293T cells further support the notion that the BAR domain of PICK1 mediates its co-aggregation with KIBRA across various cellular systems, including neurons.

### The PICK1 BAR domain regulates the interaction between PICK1 and KIBRA, as well as cohesion of the KIBRA-PICK1-GluA2 complex

Our data indicate that the PICK1 BAR domain is required for PICK1 to cluster with KIBRA. The PICK1 BAR domain is a coiled-coil structure (Fig. 4G) that can mediate protein-protein in addition to protein-lipid interactions (54, 57–59, 61, 70, 76). Additionally, the PICK1 PDZ domain can influence BAR domain interactions (56, 58, 61, 77). We observe that mutation of the PDZ domain decreases PICK1-KIBRA clustering (Fig. 4). To determine if disrupted clustering reflects decreased PICK1-KIBRA interaction, we transfected mCherry-KIBRA along with mGFP-PICK1 WT, KD-AA, 5K-E, or BAR into HEK293T cells. Immunoprecipitation of mCherry-KIBRA was performed using an mCherry antibody, and the co-immunoprecipitated PICK1 was detected using a GFP antibody (Fig. 4A). The PICK1 BAR domain and PDZ domain mutants (5K-E, KD-AA) exhibited reduced interaction with KIBRA, while the isolated PICK1 BAR domain maintained strong interaction with KIBRA (Fig. 4B). These findings indicate that the PICK1 BAR domain mediates co-clustering of PICK1 with KIBRA by influencing their interaction.

As our data demonstrate that PICK1 mediates the interaction between KIBRA and GluA2, we sought to determine the role of PICK1 BAR domain interactions in the KIBRA-PICK1-GluA2 complex. Consistent with prior work demonstrating PDZ domain/PDZ ligand-mediated interaction of PICK1 with GluA2 (37, 64), PICK1 BAR domain mutation (5K-E) does not disrupt the interaction between PICK1 and GluA2 (Fig. 4C). Subsequently, mGFP-PICK1 WT or 5K-E was co-transfected with Myc-GluA2 and mCherry-KIBRA WT into HEK293T cells to investigate how the PICK1 BAR domain mutation influences the KIBRA-PICK1-GluA2 complex (Fig. 4D). As expected, the PICK1 5K-E mutation still disrupted PICK1-KIBRA interaction in the presence of GluA2 (Fig. 4E). Surprisingly, when KIBRA was present, the PICK1 5K-E mutation also disrupted PICK1 interaction with GluA2 (Fig. 4F). These data suggest that the presence of KIBRA can alter the nature of PICK1-GluA2 complex formation, consistent with results from our clustering analysis (Fig. 2).

### Coiled-coil domains mediate KIBRA-induced protein clustering

To assess the functional domains involved in KIBRA localization and cluster formation, we transfected HEK293T cells with mGFP-KIBRA WT or various KIBRA truncations: ΔWW, ΔCoiled-coil linker (ΔCCL), ΔC2, ΔC terminal (ΔCt), or ΔαBR, Fig. 5A) along with mCherry-PICK1 WT. As previously demonstrated, KIBRA WT formed large clusters, and PICK1 co-clustered with KIBRA WT (Fig. 5B, see also Fig. 2, Fig, 3). Deletion of KIBRA’s coiled-coil linker or the C-terminal region which also contains coiled-coil structure (see Fig. 6F) resulted in diffuse distribution of KIBRA and PICK1 (Fig. 5C, 5D). Conversely, KIBRA ΔWW, ΔC2, and ΔαBR still retained the ability to form clusters, and PICK1 exhibited co-clustering with these truncations (Fig. S1). Quantification of ΔCCL or ΔCt truncations revealed a dramatic decrease in KIBRA puncta density (Fig. 5E). Both the relative PICK1 puncta density and the density of puncta co-expressing KIBRA and PICK1 also decreased with KIBRA coiled-coil linker or C-terminal truncations (Fig. 5F, G). Notably, coiled-coil domains are a common feature among diverse proteins that orchestrate biomolecular condensates through liquid-liquid phase separation (LLPS) (68, 78, 79). Indeed, recent work demonstrates that KIBRA can undergo coiled-coil domain-mediated LLPS to activate Hippo signaling (68, 80). Because the coiled-coil linker and C-terminal domain encompass the majority of KIBRA coiled-coil structure (Fig. 6F), our results suggest that the coiled-coil domains of KIBRA can mediate LLPS-like aggregation of KIBRA and PICK1.

To test if the cluster-promoting role of KIBRA’s coiled-coil-containing domains was conserved in neurons, we expressed KIBRA WT, ΔCCL, or ΔCt with PICK1 WT in hippocampal neurons. Cultured neurons were prepared from KIBRA knockout mice (14) to circumvent complications from endogenous WT KIBRA complexing with exogenously-expressed mutants. Consistent with the results in HEK293T cells, PICK1 WT formed co-clusters with KIBRA WT (Figure 5H) whereas KIBRA ΔCCL, or ΔCt showed diffuse localization and failed to cluster PICK1 (Figure 5I, J). Quantification showed a dramatic decrease in relative KIBRA puncta density (Fig. 5K), PICK1 puncta density (Fig. 5L), and KIBRA-PICK1 co-clusters (Fig. 5M) with KIBRA ΔCCL, or ΔCt mutants. These similar results between cultured neurons and HEK293T cells indicate that role of KIRBA coiled-coil domains in promoting KIBRA-PICK1 clustering are consistent across different cellular systems and exist in neurons.

### No single domain of KIBRA is responsible for mediating the interaction between KIBRA and PICK1

To identify the KIBRA domain(s) mediating interaction with PICK1, and to determine if the role of KIBRA coiled-coil-containing domains in promoting KIBRA-PICK1 clustering reflects coiled-coil-mediated protein interaction, we transfected HEK293T cells with KIBRA WT or domain deletions (ΔWW, Δcoiled-coil linker, ΔC2, ΔC-terminal, or ΔαBR) along with mCherry-PICK1 WT. PICK1 was immunoprecipitated using an mCherry antibody, and the co-immunoprecipitated KIBRA was detected with a GFP antibody (Figure 6A, B). Quantification of 3-4 biological replicates revealed that none of the KIBRA domain truncations disrupted interaction between KIBRA and PICK1 (Fig. 6C). In fact, WW domain deletion increased PICK1-KIBRA interaction (Fig. 6C), consistent with recent work demonstrating that mutation of KIBRA WW domains disrupts interaction with HIPPO signaling molecules, releasing KIBRA to increase its association with AMPA receptor complexes (50).

Multimerization plays an important role in the ability of scaffolding proteins to organize signaling complexes (68, 78, 81, 82). KIBRA can form homodimers (21, 83) and large multi-molecular clusters (Fig.2,3,5) (14, 68, 80). Yeast two-hybrid experiments indicate that KIBRA can dimerize through interactions between the CCL and C2 domains (21), whereas our data demonstrate a clear role for the CCL but not the C2 domain in mediating large, LLPS-like KIBRA cluster formation (Fig. 5, Fig. S1). To identify the modality of KIBRA homodimerization, we transfected HEK293T cells with mGFP-tagged and Myc-tagged KIBRA WT, ΔCCL, ΔC2, ΔCt, or ΔαBR (Fig. 6D). Myc-tagged KIBRA was immunoprecipitated using a Myc antibody, and the co-immunoprecipitated mGFP-tagged KIBRA counterpart was detected with a GFP antibody. Quantification of these experiments indicated that no single domain truncation disrupted KIBRA dimerization (Fig. 6E). These data suggest that KIBRA homodimerization can occur through a multimodal interaction, whereas the presence of both coiled-coil-containing domains is required for higher-order complex formation (Fig. 5).

## Discussion

### KIBRA in AMPA receptor complexes

KIBRA deletion impairs AMPA receptor trafficking, synaptic plasticity, and memory (14–16). Decreased KIBRA in the brain correlates with tauopathy-related cognitive impairment in humans, and acetylated Tau disrupts KIBRA-mediated AMPA receptor trafficking (28, 29). Conversely, increased abundance of KIBRA in AMPAR complexes has been shown to enhance hippocampal-dependent learning and memory (50). Despite these findings indicating the importance of KIBRA-dependent signaling in synaptic and brain function, the mechanism by which KIBRA organizes and regulates AMPA receptor complexes has remained elusive. The AMPAR-binding protein PICK1 was previously shown to interact with KIBRA (14), but whether and how PICK1 affected KIBRA-dependent AMPAR regulation was unknown. Here, we show that KIBRA does not interact directly with the AMPAR subunit GluA2, but that PICK1 can mediate complex formation between KIBRA and GluA2.

The large KIBRA clusters observed in our study are reminiscent of biomolecular condensates generated by LLPS (67). Indeed, recent studies demonstrated that KIBRA promotes LLPS to regulate HIPPO pathway signaling (68, 80). LLPS-organized synaptic nanodomains have emerged as mechanisms used to organize synaptic structure, segregate synaptic proteins into functional subdomains, and regulate synaptic plasticity (67, 78, 84). Together with our findings, these data raise the intriguing possibility that KIBRA may facilitate synaptic organization, signaling, and plasticity in part through promoting LLPS in excitatory postsynaptic compartments. It is important to note that KIBRA can interact with other AMPAR-binding proteins in addition to PICK1 (17, 50). Thus, the combinatorial composition of AMPAR-binding proteins recruited into KIBRA complexes may specify distinct AMPAR regulatory functions; future investigation of activity-dependent and localization-selective changes in the synaptic KIBRA interactome should provide important insight into molecular mechanisms underlying adaptive cognition.

### PICK1-KIBRA Interaction

KIBRA and PICK1 are multi-domain proteins, with each domain serving distinct functions (56, 85). Our data indicate that the PICK1 BAR domain is a crucial mediator of PICK1-KIBRA interaction, as KIBRA binds to the isolated PICK1 BAR domain and BAR domain point mutants decrease PICK1-KIBRA interaction. Supporting this co-immunoprecipitation data, isolated PICK1 BAR domain is recruited to KIBRA clusters in both HEK293T cells and neurons, whereas BAR domain mutants fail to undergo KIBRA-mediated clustering. Interestingly, mutation of the PICK1 PDZ domain also disrupts PICK1-KIBRA interaction and KIBRA-induced PICK1 clustering. This result could suggest that PICK1 PDZ domain/KIBRA PDZ ligand interactions also contribute to PICK1-KIBRA interaction (86). However, we hypothesize that these data reflect the fact that the PICK1 PDZ and BAR domains can interact, promoting a ‘closed’ conformation of PICK1 that disrupts BAR domain-dependent interactions (56, 58, 61, 77). Binding PDZ ligands promotes an open conformation of PICK1, therefore PICK1 PDZ domain mutation can indirectly inhibit BAR domain-dependent interactions through biasing PICK towards a closed conformation (56, 58, 61). Supporting this interpretation, KIBRA terminates in a type III PDZ ligand, whereas PICK1 preferentially binds type I and II PDZ ligands (51), and our data show that deletion of KIBRA’s PDZ ligand does not impair PICK1-KIBRA interaction or co-clustering. BAR domains are a specialized subtype of coiled-coil domain (60, 87). We hypothesize that heterotypic coiled-coil interactions between the PICK1 BAR and KIBRA coiled-coil domains are the primary mediators of PICK1-KIBRA interaction, and that either the coiled-coil linker or C-terminal coiled-coil regions of KIBRA are sufficient for PICK1-KIBRA interaction. Our data are consistent with this hypothesis, however, as we did not identify a single domain deletion of KIBRA that disrupted PICK1-KIBRA interaction, we cannot rule out other modes of interaction.

PICK1 binds GluA2 through PICK1 PDZ domain/GluA2 PDZ ligand interaction (37, 64). Consistently, we show that BAR domain mutation does not disrupt PICK1-GluA2 interaction, and WT KIBRA does not affect PICK1-GluA2 co-immunoprecipitation. Surprisingly, in the presence of KIBRA, mutation of the PICK1 BAR domain decreases PICK1-GluA2 interaction. PICK1-5KE substantially decreases, but does not eliminate PICK1-KIBRA interaction. These data suggest that the contact sites or orientation of interaction between KIBRA and PICK1-5KE are altered in a manner that disrupts or occludes PICK1-AMPAR interaction.

### KIBRA multimerization

We see that KIBRA forms large clusters when overexpressed in HEK293T cells, in line with previous reports demonstrating that KIBRA promotes LLPS and the formation of large biomolecular condensates to regulate HIPPO signaling in dividing cells (68, 80). Similar to Wang *et al.* (68), we find that deleting the CCL prevents formation of LLPS-like KIBRA clusters. We further demonstrate that the WW and C2 domains, αBR, and C-terminal PDZ ligand are not required for cluster formation. Our data additionally identify a critical role for the putative coiled-coil region in the KIBRA C-terminus, suggesting that both coiled-coil-containing domains of KIBRA are required for the formation of supramolecular clusters. Our data confirm previous studies reporting that KIBRA can homodimerize (21, 83), however we find that no single domain of KIBRA is essential for KIBRA dimerization. Evaluation of KIBRA dimerization using the Yeast-Two-Hybrid system showed that the KIBRA C2 domain can interact with the CCL, and deletion of the entire N-terminus of KIBRA encompassing the WW, CCL, and C2-domains prevented interaction with full-length KIBRA (21). Our data demonstrate that removing just the C2 domain does not impair KIBRA dimerization, suggesting that multiple intermolecular interactions (e.g. C2-CCL, heteromeric coiled-coil interactions between the CCL and C-terminal coiled-coil region) are able to support KIBRA dimerization. Notably, KIBRA multimerization, measured by the ability of KIBRA to form large supramolecular clusters, requires both the CCL and C-terminal coiled-coil-containing region, indicating that there are distinct requirements for KIBRA dimerization compared to higher-order multimerization (46). Given the emerging role of biomolecular condensates in organizing synaptic signaling (67, 78, 84), these findings have important implications for understanding the mechanisms by which the human memory and cognitive disorder-associated protein KIBRA regulates synaptic function and plasticity (14–17, 34).

This study reveals molecular interactions and mechanisms by which KIBRA can organize key synaptic signaling complexes. KIBRA gene and protein expression associate with human memory and multiple disorders of complex cognition, including Alzheimer’s disease, schizophrenia, autism spectrum, Tourette’s and bipolar disorders (1–13, 18–20, 88). Aberrant synaptic plasticity is a common feature across animal models relevant to these disorders. Therefore, understanding the mechanisms that can regulate neuronal responses to experience, such as molecular and synaptic plasticity, are fundamental for understanding and treating cognitive pathology.

## Experimental Procedures

### Animal Models

The University of Texas Southwestern Institutional Animal Care and Use Committees approved all animal protocols in this study. Mice were group housed in a climate-controlled environment on a 12-hour light/dark cycle. Food and water were provided *ad libitum*. Generation of KIBRA KO (14) and PICK1 KO (89) mice was described previously. Both lines were maintained on a C57Bl/6N (N10+) background.

### cDNA cloning

Mouse KIBRA and PICK1 genes were cloned from mouse tissue by RT-PCR and inserted into pCRII-TOPO (Invitrogen #K4600J10) or pCR-XL-TOPO vectors (Invitrogen #K475010). The genes were moved to a FUGW vector for transient expression or lentivirus virus generation. The mGFP, mCherry or Myc tag was inserted at N-terminal of KIBRA or PICK1. Truncations of KIBRA (delta WW domain, coiled-coil domain, C2 domain, C terminal and αPKC binding region) or PICK1 (BAR domain) were made with NEBuilder HiFi DNA Assembly Cloning Kit (NEB #E5520S). Mutations of PICK1 (KD-AA and 5K-E) were made with Q5® Site-Directed Mutagenesis Kit (NEB #E0554S). The constructs were purified by NucleoBond® Xtra Midi EF (MACHEREY-NAGEL). Myc-GluA2 was kindly provided by Dr. Richard Huganir.

### Human embryonic kidney 293T cell culture, transfection, and immunostaining

HEK293T cells were lysed in Trypsin-EDTA (0.25%) (Fisher Scientific # 25-200-072) at 37 °C for 3 mins to passage, and cultured in DMEM, high glucose, GlutaMAX™ Supplement (Life Technologies #10566016) supplied with 10% FBS (Fisher Scientific #26140079), along with 50 U/mL penicillin and 50 μg/mL streptomycin (Penicillin-Streptomycin, Fisher Scientific #15070063). HEK293T cells were transfected by calcium phosphate co-precipitation 36 h after cell passage and collected for biochemical or imaging experiments 36 h after transfection. For immunostaining, cells were fixed in 4% paraformaldehyde (Sigma-Aldrich #P6148-1KG) plus 4% sucrose (Sigma-Aldrich #S7903-5KG) in PBS at RT (room temperature) for 20 mins, permeabilized in 0.2% TritonX-100 in PBS at RT for 10mins, and blocked in 10% NDS (normal donkey serum, Jackson ImmunoResearch #017-000-121) in PBS at RT for 1 h. Cells were then incubated with primary Myc-Tag (9B11) Mouse mAb (Cell Signaling Technology #2276S) in PBS + 3% NDS at RT for 1 h followed by Goat anti Mouse IgG (H+L) Cross-Adsorbed Secondary Antibody (DyLight 405, Invitrogen #PI35500) in PBS + 3% NDS at RT for 1 h. After immunostaining, Cells were mounted in Fluoroshield histology mounting medium (Sigma-Aldrich #F6182-20ML) and stored at RT until imaging.

### Lentivirus production and purification

FUGW mGFP/mCherry-KIBRA (WT, delta coiled-coil domain or C terminal) or PICK1 (WT, 5K-E or BAR domain) was transfected with REV, RRE and VSV-G into HEK293T cells 36 h after cell passage. Lentivirus was produced form HEK293T cells and released to the supernatant. Supernatant containing the viruses was collected 2 days after transfection and centrifuged at 1000g for 5 min at RT to pellet any cell debris. The viral supernatant was concentrated through Amicon Ultra-15 centrifugal filters and stored at –80 °C

### Neonatal mouse hippocampal neuronal culture and transfection

Hippocampi were dissected from neonatal P0 homozygous KIBRA KO or PICK1 KO mice and incubated with 670 μg/mL Papain (Worthington #LS003119) and 100 μg/mL DNase (Sigma-Aldrich #DN-25) at 37 °C for 10 minutes. Neurons (2.5×10^5^/mL) were plated on poly-L-lysine (Sigma #P2636) coated coverslips in Neurobasal growth medium (Fisher Scientific #21103049) supplemented with 2% B27 (Fisher Scientific #17504044), 2 mM Glutamax (Fisher Scientific #35-050-061), 50 U/mL penicillin, 50 μg/mL streptomycin (Penicillin-Streptomycin, Fisher Scientific #15070063) and 5% Donor Equine Serum (Cytiva #SH30074.03). Three hours after plating, the medium was changed to Neurobasal growth medium supplemented with 2% B27, 2 mM Glutamax, along with 50 U/mL penicillin and 50 μg/mL streptomycin. Neurons were transfected with Lentivirus at DIV10 and observed at DIV14.

### Co-immunoprecipitation and western blot

HEK293T cells cultured in 60 mm dishes were lysed in 1 ml PBS with 1% TritonX-100 plus protease inhibitors (cOmplete™, Mini Protease Inhibitor Cocktail, Sigma-Aldrich #11836153001) and phosphatase inhibitors (1mM NaPPi, 5mM NaF, 1mM NaVO3). Protein concentration was assayed by Pierce Detergent Compatible Bradford Assay Kit (Thermo Fisher Scientific # 23246). 25-50 ul Protein G Magnetic Beads (NEB #S1430S) were incubated with 3ug GFP (D5.1, Rabbit mAb, Cell Signaling Technology #2956S), mCherry (E5D8F, Rabbit mAb, Cell Signaling Technology #43590S) or Myc (9B11, Mouse mAb, Cell Signaling Technology #2276S) antibody at 4 °C for 1 h. Cell lysate was incubated with antibody-conjugated beads at 4 °C overnight. The incubated beads were washed in 500ul lysis buffer once, in 500 ul lysis buffer plus 0-500 mM NaCl twice and in 500 ul PBS once. The immunoprecipitated proteins were eluted by SDS protein sample buffer (10% glycerol, 62.5mM Tris/HCl pH 6.8, 2% sodium dodecyl sulfate, 0.01% bromophenol blue, 1.25% beta-mercaptoethanol). Samples were separated via SDS-PAGE (8% gels) then transferred to nitrocellulose membrane (LI-COR #926-31092) or PVDF membrane (GE Healthcare #10600023) in cold (∼4°C) transfer buffer (25 mM Tris, 192 mM glycine, 0.03% sodium dodecyl sulfate and 20% methanol). Membranes was blocked with 2% milk for at RT for 1 h. then incubated with primary antibodies (anti-GFP, mCherry or Myc, Catalog numbers listed above) at 4°C overnight. Membranes were then washed 5 times with TBST (10 minutes per wash) followed by incubation with fluorescent secondary antibodies (anti-mouse or anti-rabbit Licor IRDye 680RD or 800CW, #926-68070, 926-68071, 926-32210 or 926-32211) or HRP-conjugated secondary antibodies (Chicken anti Mouse secondary antibody HRP, Fisher Scientific # A15981, Sheep anti Rabbit secondary antibody HRP, Fisher Scientific #A16172) at RT for 1 h, then a second round of 5x 10 minute washes. Membranes incubated with HRP-conjugated secondary antibodies were incubated with Amersham ECL Prime Western Blotting Detection Reagent (GE Healthcare #RPN2232) for 5 mins. Fluorescence and Chemiluminescence were imaged using the ChemiDoc MP Imaging System from Bio-Rad, which provides a measure of signal saturation to ensure that no bands are saturated. All westerns were normalized to input and then to control sample run on the same gel.

### Confocal imaging and data analysis

All samples were imaged with Zeiss LSM 710 Confocal Microscope at RT. mCherry, mGFP, and immnostained Myc were imaged at 561 nm, 488 nm and 405nm excitation, respectively. HEK293T cells were imaged through a 20× dry objective. Neurons were imaged through a 63× oil objective. 3D images were captured at 1 µm intervals as a single optical section. Images were analyzed using ImageJ software. For puncta density analysis, this entailed projecting image Z stacks using the Sum Slices, splitting color channels, subtracting background and thresholding for each image with the experimenter blind to experimental conditions. Colocalization was determined by overlapping different channels using Image-Overlay-Add Image function. The number of particles was quantified using the analyze particles function. Analyses in neurons were applied to soma and neurites. For line scan analyses a single optical slice was analyzed using the ‘Plot Profile’ function in Image J, determined by the brightest slice in the KIBRA and/or PICK1 channels. The lowest value of the line scan was subtracted as local background.

### Quantification and statistical analysis

Statistical analysis was performed in Graph Pad Prism version 9.4.1. Data are plotted as mean ± SD unless otherwise noted. The statistical test performed for each analysis is noted in the figure legend. p< 0.05 was considered significant. Data were quantified from 3-4 biological replicates representing distinct batches of HEK293T cells, each containing more than 100 cells, or neuronal cultures prepared from different litters, each grown on more than 1 coverslip. Statistical tests were chosen based on sample size, hypotheses, and agreement with statistical assumptions.

## Data availability

All data are contained within the manuscript.

## Author contributions

**Xin Shao:** Conceptualization, methodology, formal analysis, investigation, writing-original draft, writing-review and editing, visualization.

**Lenora Volk:** Conceptualization, formal analysis, writing-original draft, writing-review and editing, visualization, funding acquisition.

---

**Figure S1.**
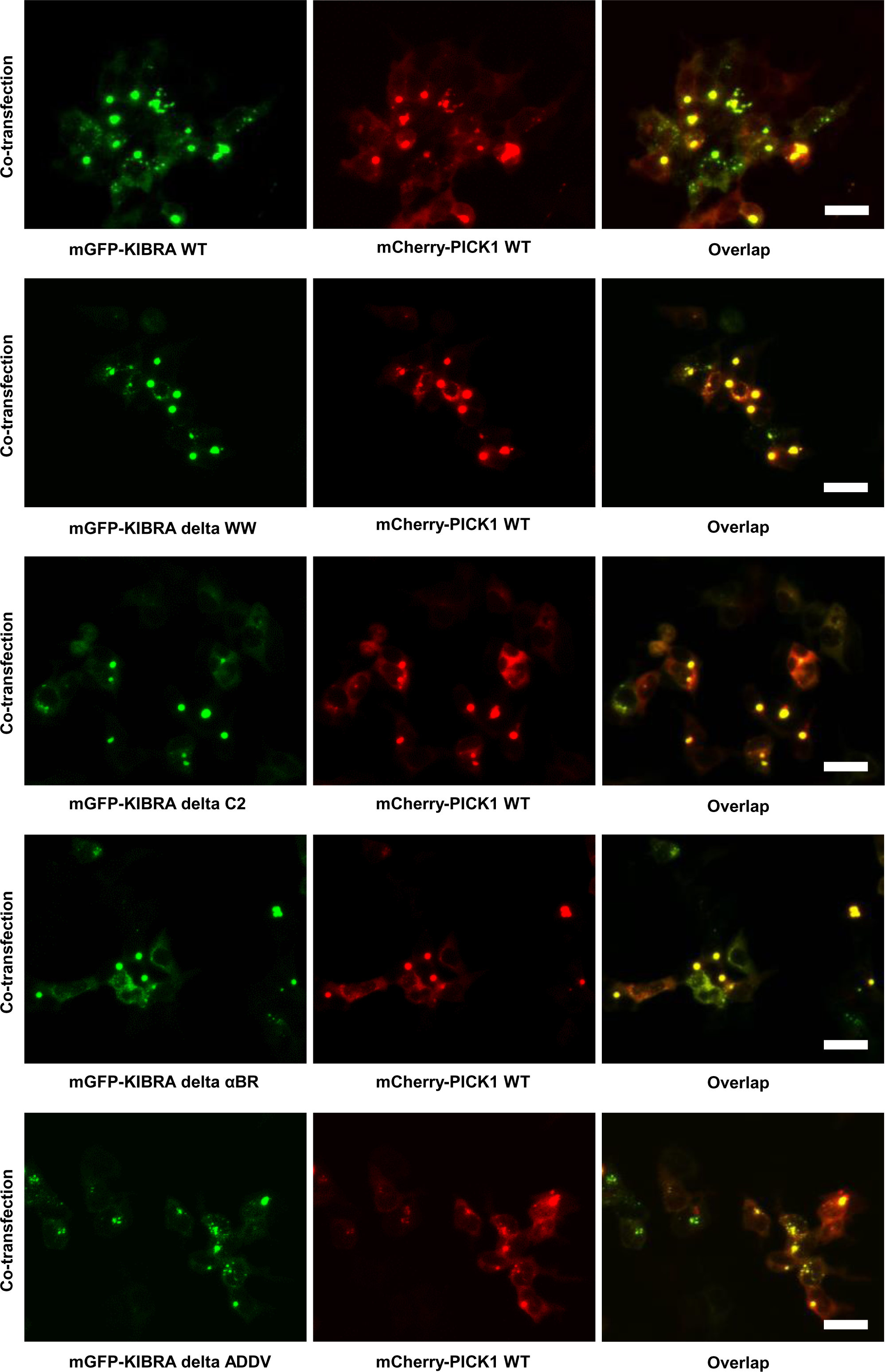
KIBRA WW and C2 domains, aPKC-binding region, and PDZ ligand are not required for KIBRA clustering or PICK1 co-aggregation with KIBRA. Overexpressed KIBRA WT and KIBRA mutants missing WW, C2, αBR or ADDV domains cluster with PICK1 in HEK293T cells. Scale bar: 30 µm.

